## Supplemental Material for "Kinematics and morphological correlates of descent strategies in arboreal mammals suggest early upright postures in euprimates"

### Supplementary material

This document includes:
Tables S1, S2, S3 and S4
Figures S1, S2, and S3

See also the additional Excel file “SuppTables” for complete results of statistical analysis.

**Table S1. List of animals studied.** Species with an* are nocturnal. PZP= Parc Zoologique de Paris, France. PZBM= Parc Zoologique et Botanique de Mulhouse, France.

| | Family | Subfamily | *Species* | Housing location | Observation period | Sample size | Year of birth |
|---|---|---|---|---|---|---|---|
| Strepsirrhines primates | Lorisidae | Lorisinae | *Xanthonycticebus pygmaeus** | Nowe Zoo Poznan, Poland | 03-2013 | 4 females | unknown |
| | | | *Hapalemur occidentalis* | Parc Zoologique de Paris (PZP), France | 04-2017 | 1 male | 2014 |
| | | | *Hapalemur griseus* | Parc Zoologique et Botanique de Mulhouse (PZBM), France | 09-2016 | 1 female | 2006 |
| | Lemuridae | | *Eulemur rubriventer* | PZBM | 09-2016 | 2 females, 2 males | 1991, 2010 & 2008, 2009 |
| | | | *Eulemur coronatus* | PZBM | 09-2016 | 1 female, 3 males | 1999 & 2005, 2015, 2015 |
| | | | *Eulemur mongoz* | PZBM | 09-2016 | 1 female | 1987 |
| | | | | PZP | 04-2017 | 2 males | 1996, 2001 |
| Platyrrhines primates | Pitheciidae | Callicebinae | *Plecturocebus cupreus* | PZP | 04-2017 | 1 female, 1 male | 2007 & 2010 |
| | | Saimirinae | *Saimiri boliviensis boliviensis* | PZBM | 01-2016 | 2 females, 2 males | 2005, 2010 & 2011, 2015 |
| | Cebidae | Aotinae | *Aotus lemurinus griseimembra** | Spaycific'Zoo, France | 09-2017 | 1 female, 1 male | 2010 & 2009 |
| | | Callitrichinae | *Saguinus imperator* | PZBM | 09-2016 | 2 females, 2 males | 2010, 2013 & 2014, 2014 |
| | | | *Saguinus oedipus* | Spaycific'Zoo, France | 09-2017 | 1 female, 1 male | 2016 & 2015 |
| Scandentians | Tupaiidae | | *Tupaia belangeri* | Moscow Zoo, Russia | 07-2017 | 3 females | unknown |
| Rodents | Gliridae | Leithiinae | *Dryomys nitedula** | University of Thessaloniki, Greece - wildcaught | 07-2017 | 1 female, 1 male | unknown |
| | | Graphiurinae | *Graphiurus murinus** | Moscow Zoo, Russia | 07-2017 | 1 female, 1 male | unknown |
| | Platacanthomyidae | | *Typhlomys chapensis** | Moscow Zoo, Russia | 07-2017 | 2 males | unknown |
| Carnivorans | Procyonidae | | *Procyon lotor* | Spaycific'Zoo, France | 09-2017 | 1 female, 3 males | 2005 & 2007, 2012, 2012 |
| | | | *Nasua nasua* | Spaycific'Zoo, France | 09-2017 | 2 females, 2 males | unknown |
| | | | *Potos flavus** | Spaycific'Zoo, France | 09-2017 | 1 female | unknown |
| Marsupials | Phalangeridae | Caluromyinae | *Caluromys philander* | Laboratoire d'écologie générale de Brunoy, France | 03-1993 | 2 males | 1990, 1991 |
| | | | *Trichosurus vulpecula** | Spaycific'Zoo, France | 09-2017 | 1 female, 1 male | unknown |
| | Petauridae | | *Petaurus breviceps** | Spaycific'Zoo, France | 09-2017 | 2 females, 2 males | 2010, 2014 & 2012, 2014 |

**Table S2. Mean body mass, endocranial volume (ECV) and encephalization quotient (EQ) used for each extant and extinct species studied and their associated references.** See Table S3 for the details of ECV measurement conducted in this study.

| **Extant species** | | | | | | |
|---|---|---|---|---|---|---|
| | **Species** | **Mean body mass (g)** | **Reference for body mass** | **ECV (mL)** | **Calculated EQ** | **Reference for ECV** |
| Strepsirrhines | *Xanthonycticebus pygmaeus* | 362.5 | (Fleagle, 2013) | 7.23 | 1.6484 | (Powell et al., 2017) |
| | *Hapalemur occidentalis* | 1017 | Fleagle 2013 | 13.75 | 1.4522 | Powell et al. 2017 |
| | *Hapalemur griseus* | 709 | Fleagle 2013 | 13.78 | 1.9048 | Powell et al. 2017 |
| | *Eulemur rubriventer* | 1960 | Fleagle 2013 | 23.91 | 1.5478 | Powell et al. 2017 |
| | *Eulemur coronatus* | 1180 | Fleagle 2013 | 19.78 | 1.8697 | Powell et al. 2017 |
| | *Eulemur mongoz* | 1212.5 | Fleagle 2013 | 18.89 | 1.7498 | Powell et al. 2017 |
| Platyrrhines | *Plecturocebus cupreus* | 1070 | Fleagle 2013 | 17.44 | 1.7734 | Powell et al. 2017 |
| | *Saimiri boliviensis boliviensis* | 871.5 | Fleagle 2013 | 21.52 | 2.5502 | Powell et al. 2017 |
| | *Aotus lemurinus griseimembra* | 966 | Fleagle 2013 | 16.07 | 1.7636 | Powell et al. 2017 |
| | *Saguinus imperator* | 474.5 | Fleagle 2013 | 10.7 | 1.9956 | Powell et al. 2017 |
| | *Saguinus oedipus* | 411 | Fleagle 2013 | 9.7 | 2.0138 | Powell et al. 2017 |
| Scandentia | *Tupaia belangeri* | 160 | animaldiversity.org (Myers et al., 2021) | 2.15 | 0.9023 | This study |
| Rodents | *Dryomys nitedula* | 26 | animaldiversity.org | 0.58 | 0.9441 | This study |
| | *Graphiurus murinus* | 28.5 | animaldiversity.org | 0.46 | 0.6992 | This study |
| | *Typhlomys chapensis* | 16.93 | (Cheng et al., 2017) | 0.4 | 0.8967 | This study |
| Carnivorans | *Procyon lotor* | 6000 | animaldiversity.org | 40.04 | 1.1250 | (Finarelli, 2006) |
| | *Nasua nasua* | 4500 | animaldiversity.org | 29.96 | 1.1609 | Finarelli et al. 2006 |
| | *Potos flavus* | 3300 | animaldiversity.org | 25.53 | 1.1205 | Finarelli et al. 2006 |
| Marsupials | *Caluromys philander* | 265 | animaldiversity.org | 3.5 | 1.0081 | (Ashwell, 2008) |
| | *Trichosurus vulpecula* | 2850 | animaldiversity.org | 11 | 0.5386 | Ashwell et al. 2008 |
| | *Petaurus breviceps* | 110 | animaldiversity.org | 1.3 | 0.7215 | This study |

| **Extinct species** | | | | | |
|---|---|---|---|---|---|
| | ***Species*** | **Mean body mass (g)** | **Mean EQ** | **Estimated age (MY)** | **Reference for EQ, body mass and pictures for body lengths measured** |
| Adapiformes | *Darwinius masillae* | 660 | NA | 47 | (Franzen et al., 2009) |
| | *Europolemur kelleri* | 1485 | NA | 45 | (Koenigswald, 1979) |
| | *Notharctus tenebrosus* | 4947 | 0.26 | 45 | (Harrington et al., 2016) |
| | *Adapis parisiensis* | 1544 | 0.56 | 35 | (Ramdarshan and Orliac, 2016) |
| Omomyiformes | *Archicebus achilles* | 25 | NA | 55 | (Ni et al., 2013; Dagosto et al., 2018) |
| | *Microchoerus erinaceus* | 642 | 0.53 | 35 | (Ramdarshan and Orliac, 2016) |
| | *Tetonius homunculus* | 113 | 0.7 | 50 | (Ramdarshan and Orliac, 2016) |

| | | | | | |
|---|---|---|---|---|---|
| Plesiadapiformes | *Plesiadapis tricuspidens* | 2039 | 0.26 | 60 | (Orliac et al., 2014) |
| | *Plesiadapis cookei* | 2200 | 0.31 | 60 | (Silcox et al., 2009; Bertrand and Silcox, 2016) |
| | *Plesiadapis insignis* | 783 | NA | 60 | (Boyer, 2009) |
| | *Carpolestes simpsoni* | 100 | NA | 58 | (Silcox et al., 2017) |
| | *Ignacius graybullianus* | 375 | 0.48 | 58 | (Silcox et al., 2009; Harrington et al., 2016) |
| Rodent | *Ischyromys typus* | 1342.23 | 0.47 | 30 | (Bertrand and Silcox, 2016) |

**Table S3. Endocranial volume (ECV) measured on specimens of various extant species.** Specimens are from the anatomical collections of the Museum für Naturkunde of Berlin, Germany. We used Chia seeds, with a calculated reference of 1g = 1.22 ml. We calculated the EQ using a reference brain volumic mass of 1.036 g.cm$^{-3}$ (Ebinger, 1974) and the following formula (Boddy et al., 2012):

$$EQ = \frac{1.036 \times ECV}{0.056 \times BodyMass^{0.746}}$$

| Species | Specimen number | Calculated ECV (ml) | Mean ECV (ml) | STD ECV |
|---|---|---|---|---|
| *Tupaia belangeri* | ZMB_Mam_90819 | 2.2448 | 2.15126667 | 0.14140075 |
| *Tupaia belangeri* | ZMB_Mam_90780 | 1.9886 | | |
| *Tupaia belangeri* | ZMB_Mam_90781 | 2.2204 | | |
| *Dryomys nitedula* | ZMB_Mam_I1062 | 0.5978 | 0.57746667 | 0.01863581 |
| *Dryomys nitedula* | ZMB_Mam_I1065 | 0.5612 | | |
| *Dryomys nitedula* | ZMB_Mam_I1085 | 0.5734 | | |
| *Graphiurus murinus* | N.A. | 0.5368 | 0.45953333 | 0.07937817 |
| *Graphiurus murinus* | ZMB_Mam_20339 | 0.3782 | | |
| *Graphiurus murinus* | ZMB_Mam_45008 | 0.4636 | | |
| *Typhlomys cinereus* | ZMB_Mam_16777 | 0.4026 | 0.4026 | NA |
| *Petaurus breviceps* | ZMB_Mam_86756 | 1.403 | 1.2993 | 0.14665395 |
| *Petaurus breviceps* | ZMB_Mam_3471 | 1.1956 | | |

NA= not applicable

**Table S4. Calculated limbs proportions for the studied extinct species.** See Table S2 for the associated mean body masses, mean EQ and references from where we extracted the photographs used to measure the postcranial lengths.

| Species | % Tail to body length | % Forelimb to body length | % Hindlimb to body length | Intermembral index | % Hand to forelimb length | % Foot to hindlimb length | % Pollex to hand length | % Hallux to foot length |
|---|---|---|---|---|---|---|---|---|
| *Archicebus achilles* | NA | NA | NA | NA | NA | **59.65** | NA | **50** |
| *Microchoerus erinaceus* | NA | NA | NA | NA | NA | NA | NA | NA |
| *Tetonius homunculus* | NA | NA | NA | NA | NA | NA | NA | NA |
| *Adapis parisiensis* | NA | NA | NA | NA | NA | NA | NA | NA |
| *Darwinius masillae* | **131.71** | **49.59** | **80.89** | **59.40** | **54.43** | **49.62** | **51.16** | **46.97** |
| *Europolemur kelleri* | NA | NA | NA | NA | NA | **50.29** | NA | **40.23** |
| *Notharctus tenebrosus* | NA | NA | NA | NA | NA | NA | NA | NA |
| *Plesiadapis cookei* | **107.69** | **54.62** | **64.62** | **95.54** | **42** | **60.51** | NA | NA |
| *Plesiadapis insignis* | **68.38** | **54.27** | **59.83** | **92.47** | **47.67** | **50.54** | NA | **34.04** |
| *Plesiadapis tricuspidens* | NA | NA | NA | NA | NA | NA | NA | NA |
| *Carpolestes simpsoni* | NA | **50.36** | **63.31** | **78.33** | **48.94** | **46.67** | NA | **39.29** |
| *Ignacius graybullianus* | NA | NA | NA | NA | NA | NA | NA | NA |
| *Ischyromys typus* | NA | NA | NA | NA | NA | NA | NA | NA |

NA= not applicable

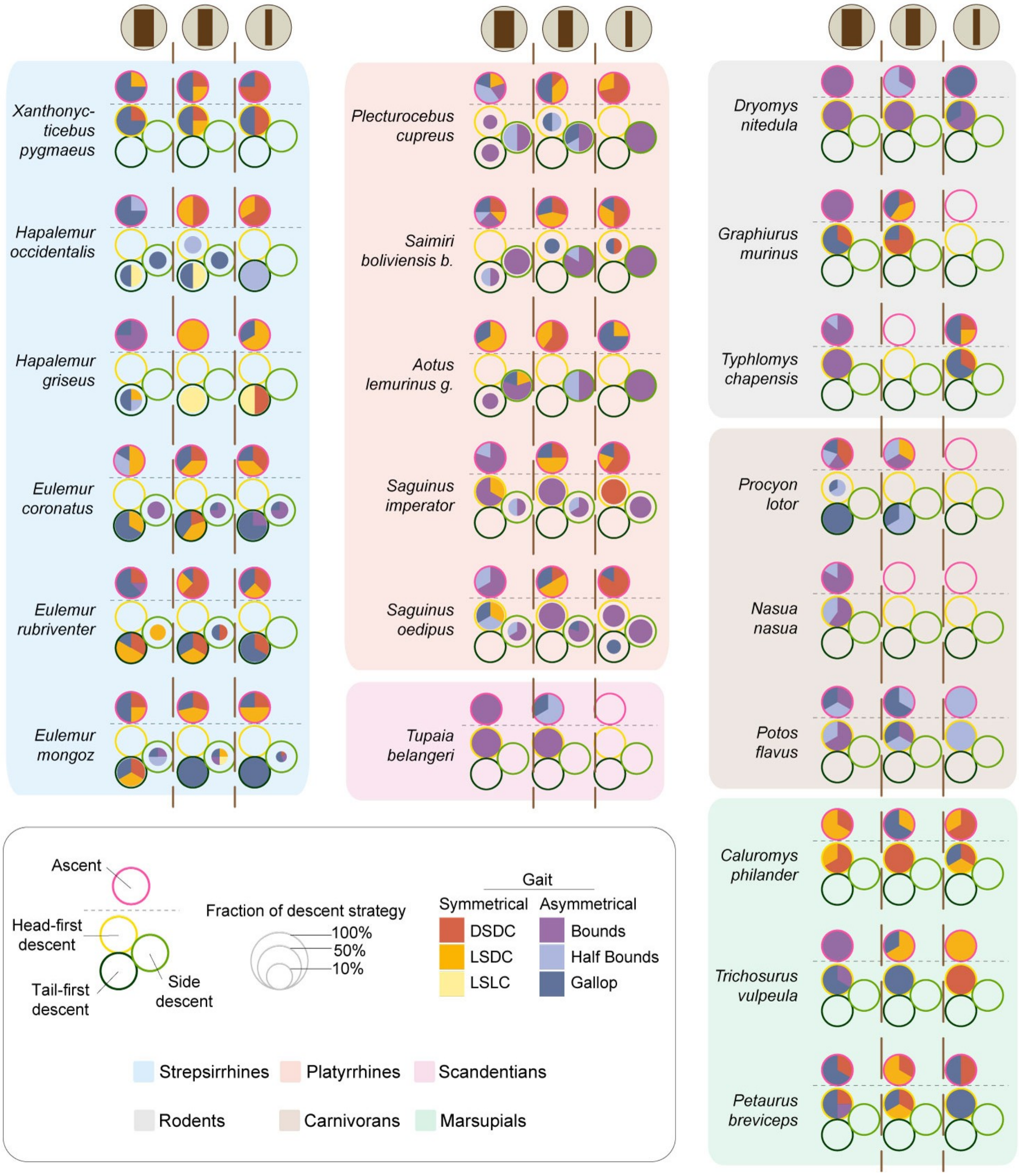


**Figure S1. Mean proportions of gait types (in %) by support diameter and by ascent and descent conditions, for each species.** DSDC: diagonal sequence – diagonal couplet, LSDC: lateral sequence – diagonal couplet, LSLC: lateral sequence – lateral couplet.

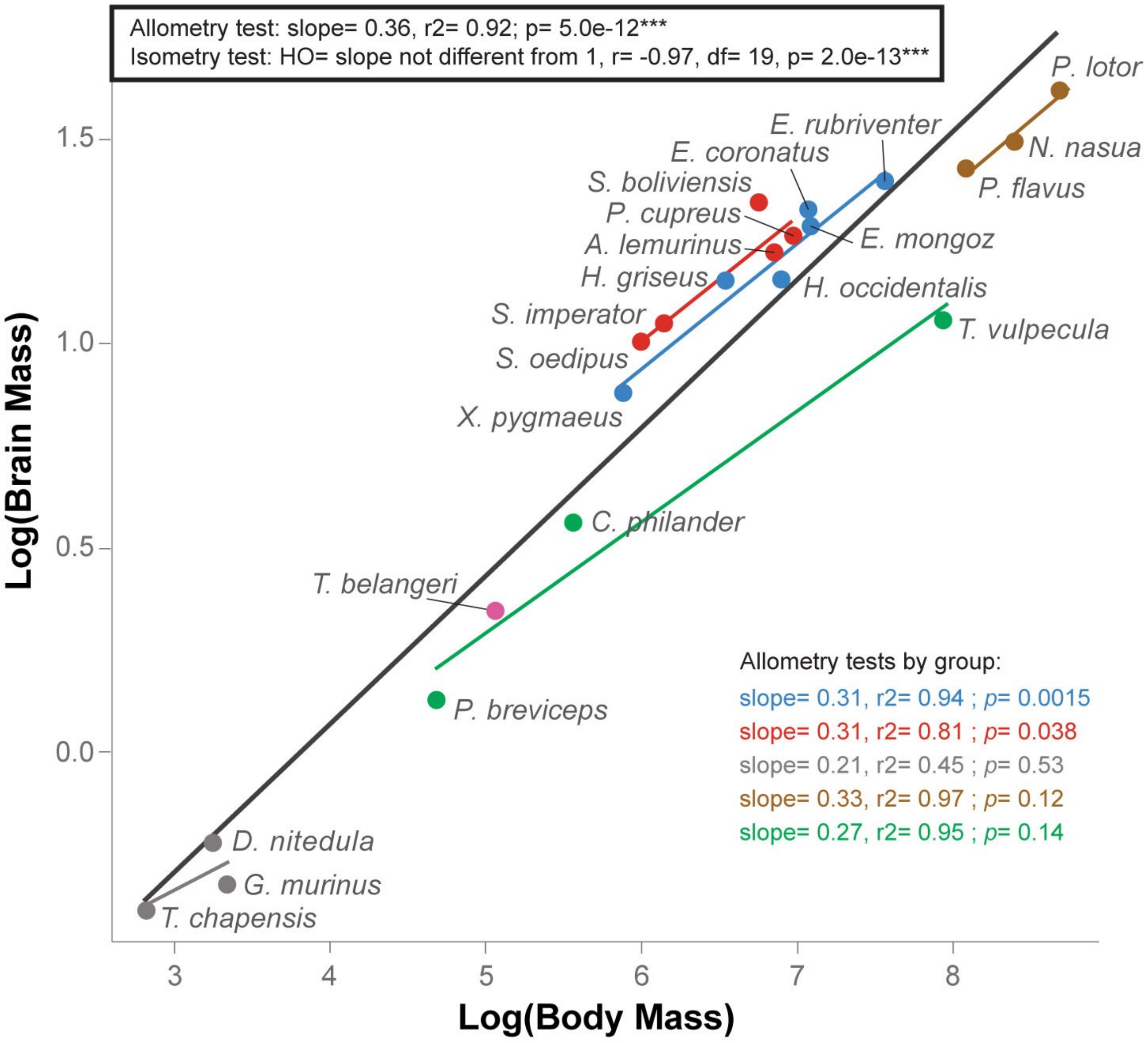


**Figure S2. Major axis orthogonal regressions on the Log transformed mean brain and mean body masses for all species (in black with the grey area representing the associated confidence interval) and by phylogenetic groups color coded as defined in Fig. 1C.** Brain masses were calculated from ECV using a reference brain volumic mass of 1.036 g.cm3 (Ebinger, 1974). See Table S2 for associated mean body masses and mean endocranial volumes (ECV) by species.

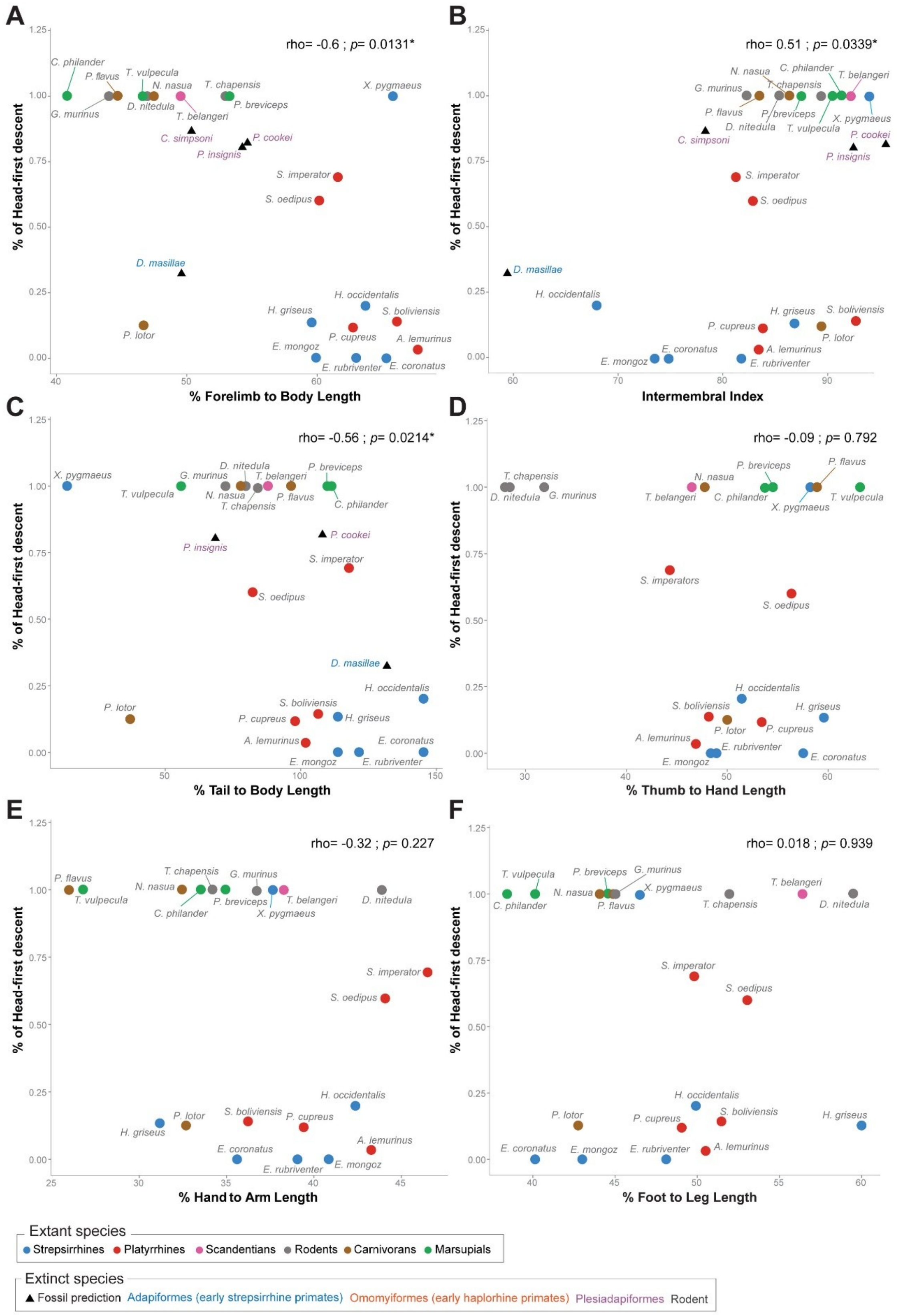


**Figure S3.** Spearman correlations and associated corrected p-values between the proportion of head-first descents by species on vertical supports (all diameters combined) and the relative: **A.** forelimb length, **B.** intermembral index, **C.** tail length, **D.** thumb length, **E.** hand length, and **F.** foot length. See Fig. 4 for the correlations on other variables, Table 1 for the variables list and descriptions, and SuppTables G and H for spearman rhos and associated corrected p-values of all variables.
